# Spatial landscape of malignant pleural and peritoneal mesothelioma tumor immune microenvironment

**DOI:** 10.1101/2023.09.06.556559

**Authors:** Xiaojun Ma, David Lembersky, Elena S Kim, Tullia C Bruno, Joseph R. Testa, Hatice U Osmanbeyoglu

## Abstract

Immunotherapies have shown modest clinical benefit thus far for malignant mesothelioma (MM). A deeper understanding of immune cell spatial distribution within the tumor immune microenvironment (TIME) is needed to identify interactions between tumor and different immune cell types that might impact the effectiveness of potential immunotherapies. We performed multiplex immunofluorescence (mIF) using tissue microarrays (TMAs, n=3) of samples from patients with malignant peritoneal (n=25) and pleural (n=88) mesothelioma (MPeM and MPM, respectively) to elucidate the spatial distributions of major immune cell populations and their association with LAG3, BAP1, NF2, and MTAP expression, the latter as a proxy for CDKN2A/B. We also analyzed the relationship between the spatial distribution of major immune cell types with MM patient prognosis and clinical features. The distribution of immune cells within the TIME is similar between MPM and MPeM. However, there is a higher level of interaction between immune cells and tumor cells in MPM than MPeM. Within MPM tumors, there is increased amount of interaction between tumor cells and CD8^+^ T cells in BAP1-low than in BAP1-high expressing tumors. The cell-cell interactions identified in this investigation have potential implications for the immune response against MM tumors and could be a factor in the different behaviors of MPM and MPeM. Our findings provide a valuable resource for the MM cancer research community and exemplifies the utility of spatial resolution within single-cell analyses.

Our mesothelioma spatial atlas mIF dataset is available at https://mesotheliomaspatialatlas.streamlit.app/.

## Introduction

Malignant mesothelioma (MM) is an uncommon but highly aggressive cancer of serosal surfaces linked to asbestos exposure^1,2^. MM primarily affects the lining of the lungs (pleural), but it can also occur in the linings of the abdomen (peritoneal), heart (pericardial), or tunica vaginalis covering the testicles (testicular). Malignant pleural mesothelioma (MPM) comprises 80% of cases, while malignant peritoneal mesothelioma (MPeM) includes 15-20% of cases^3^. MM is broadly divided into three main histological subtypes with varying biological and clinical behaviors: epithelioid (most common type), sarcomatoid and biphasic, the latter being a combination of the former types. MPeM has a significantly better prognosis than MPM, with a median overall survival exceeding 5 years when amenable to cytoreductive surgery and hyperthermic intraperitoneal chemotherapy^3^. The prognosis for MPM remains poor, with a 5-year survival rate of <10%^4^. Men are at a higher risk than women, likely due to higher occupational exposure. However, MM is increasingly being diagnosed in younger individuals, and in women, with no known history of asbestos exposure^5^.

There is no cure for MM. Currently approved frontline therapies for MPM, including immunotherapy (nivolumab plus ipilimumab) and chemotherapy (cisplatin plus pemetrexed), modestly extend the overall survival by only a few months^6,7^. This poor outcome may be influenced by the complex structure of MM tumor microenvironment. Recent exploratory biomarker analyses showed a significantly longer median overall survival (OS) among immunotherapy-treated MPM patients with an inflammatory gene signature score (based on levels of CD8A, CD274/PD-L1, STAT1, and LAG3) regardless of histology^8^.

Over the past few decades, there has been significant progress in understanding the molecular mechanisms underlying MM development and progression. MPM and MPeM have similarities of cellular features and common genetic alterations. The genomic profile of MM reveals a low protein-coding mutation rate, with the most commonly occurring alterations being mutations and deletions of the tumor suppressor genes (TSGs) BRCA1 associated protein-1 (*BAP1*; located at chromosome band 3p21), neurofibromatosis type 2 (*NF2*; 22q12), and cyclin-dependent kinase inhibitor 2A and 2B (*CDKN2A/B*; 9p21)^9-14^. Recently, we provided a comprehensive analysis of data on MPM tumors with genomic alteration in one or more of these specific TSGs, which are thought to be critical drivers of MPM pathogenesis^15^. Although alterations of these key TSGs frequently occur in various combinations in a given MM tumor, we showed that alteration of *BAP1* alone is associated with a better clinical outcome and an immunotherapy response signature. Interestingly, *BAP1* alterations have been shown recently to be correlated with perturbed immune signaling in MPeM^16^. Furthermore, MPM tumors with alterations in *BAP1* alone showed a distinct pattern of expression of inflammatory tumor microenvironment genes, including activation of interferon signaling and IRF transcription factors (TFs) and high *LAG3* (lymphocyte activation gene-3) and *VISTA* (V-domain Ig suppressor of T cell activation) immune checkpoint expression^15^. *LAG3* is expressed on activated T cells and has now become a part of the repertoire of combinatorial immunotherapies available for the treatment of metastatic melanoma^17^.

A deeper understanding tumor immune microenvironment (TIME) and associated molecular and clinical features, which differentiate or associate the two major MM types is crucial for the development of effective treatment strategies. Spatial technologies have revealed the complexity of the TIME with unparalleled resolution and have increasingly been shown to be of prognostic and predictive importance^4^. The aim of this study was to enhance our understanding of the TIME along with the protein expression of LAG3, BAP1, NF2, and MTAP (with MTAP acting as a surrogate for CDKN2A/B, since the *MTAP* gene at 9p21 is often co-deleted with *CDKN2A/B* in MM) within distinct types of MM. To achieve this, we employed multiplex immunofluorescence (mIF) techniques on TMA sections of MPM and MPeM samples.

## Results

### Mesothelioma multiplex immunofluorescence dataset and patient characteristics

To spatially characterize the cellular landscape of the MM TIME, we applied mIF to samples from patients with MPM and MPeM (**Fig. 1A**). Briefly, we obtained three TMAs consisting of 336 samples/cores of MM from a total of 115 patients from the National Mesothelioma Virtual Bank (NMVB), a database funded by the Center for Disease Control and Prevention (CDC) in association with National Institutes of Occupational Health and Safety (NIOSH). There were 1–9 cores per patient, and only 13 patients were represented with one core. These patients were seen at Roswell Park Cancer Institute (RPCI, n=53), University of Pennsylvania (UPENN, n=30), or University of Pittsburgh/UPMC (PITT, n=32). Information on cohort characteristics and associated clinicopathological features are described in **Tables 1** and **2**.

**Figure 1:**
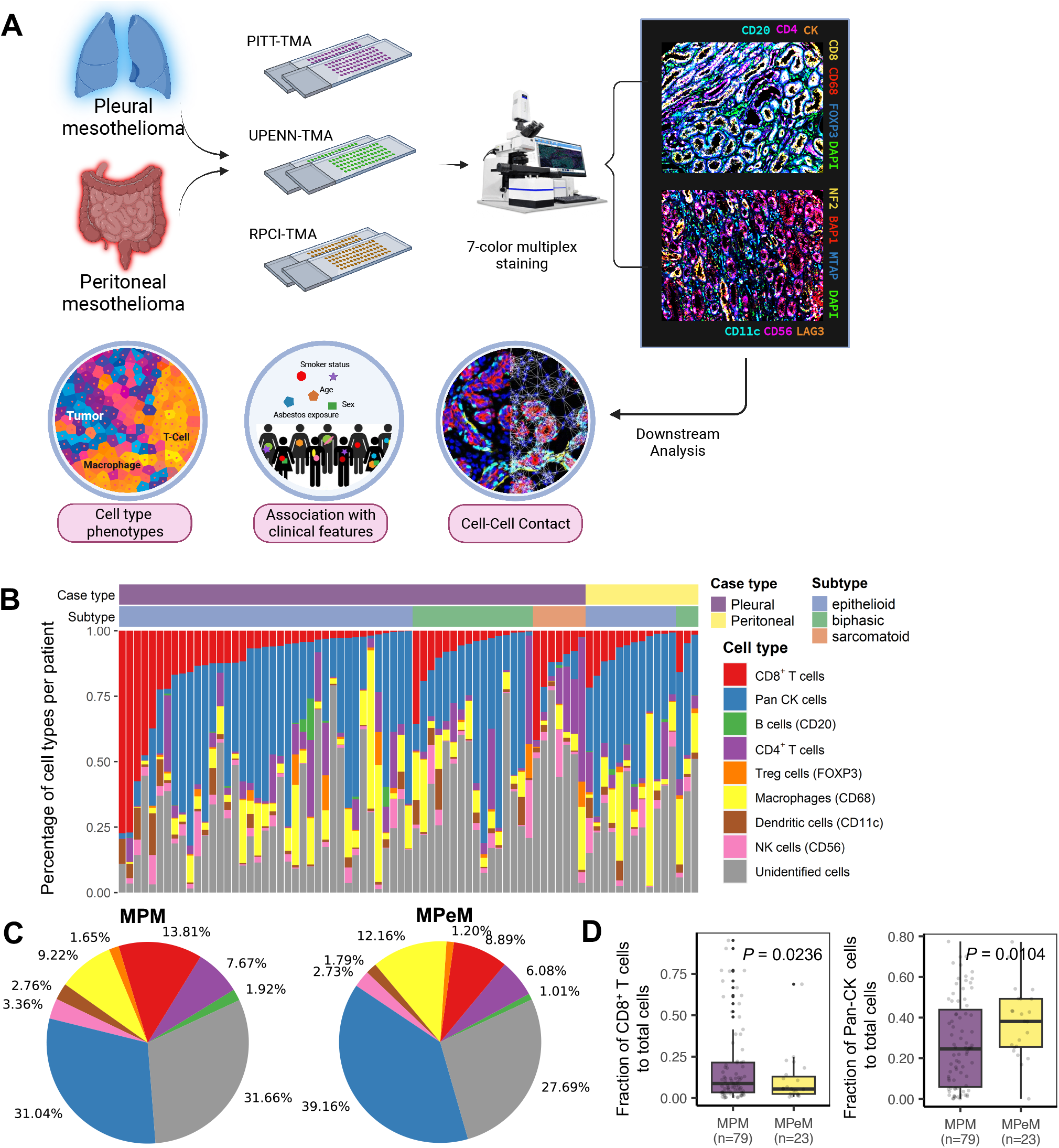
Multiplex immunofluorescence defines the spatial landscape of MM. **A)** Schematic diagram depicting acquisition of multiplexed images from MM patients. Figure was created with BioRender. **B)** Waterfall plot depicting the distribution of cell populations as a percentage of all cells in the TIME, sorted by CD8^+^ T cells across histological subgroups within MPM and MPeM. Cell percentages are displayed as vertical bars (colors correspond to cell lineages). **C)** Pie charts representing mean cell composition in MPM (n=79) and MPeM (n=23) patients including relative frequencies of B cells, CD4^+^ T cells, CD8^+^ T cells, Treg cells, macrophages, DCs, NK, Pan-CK^+^ cells and unidentified cells. **D)** Prevalence of CD8^+^ T and Pan-CK^+^ cells across MPM and MPeM as a fraction of total cells. *P*-value was calculated using the one-sided Mann–Whitney U test.

**Table 1:**
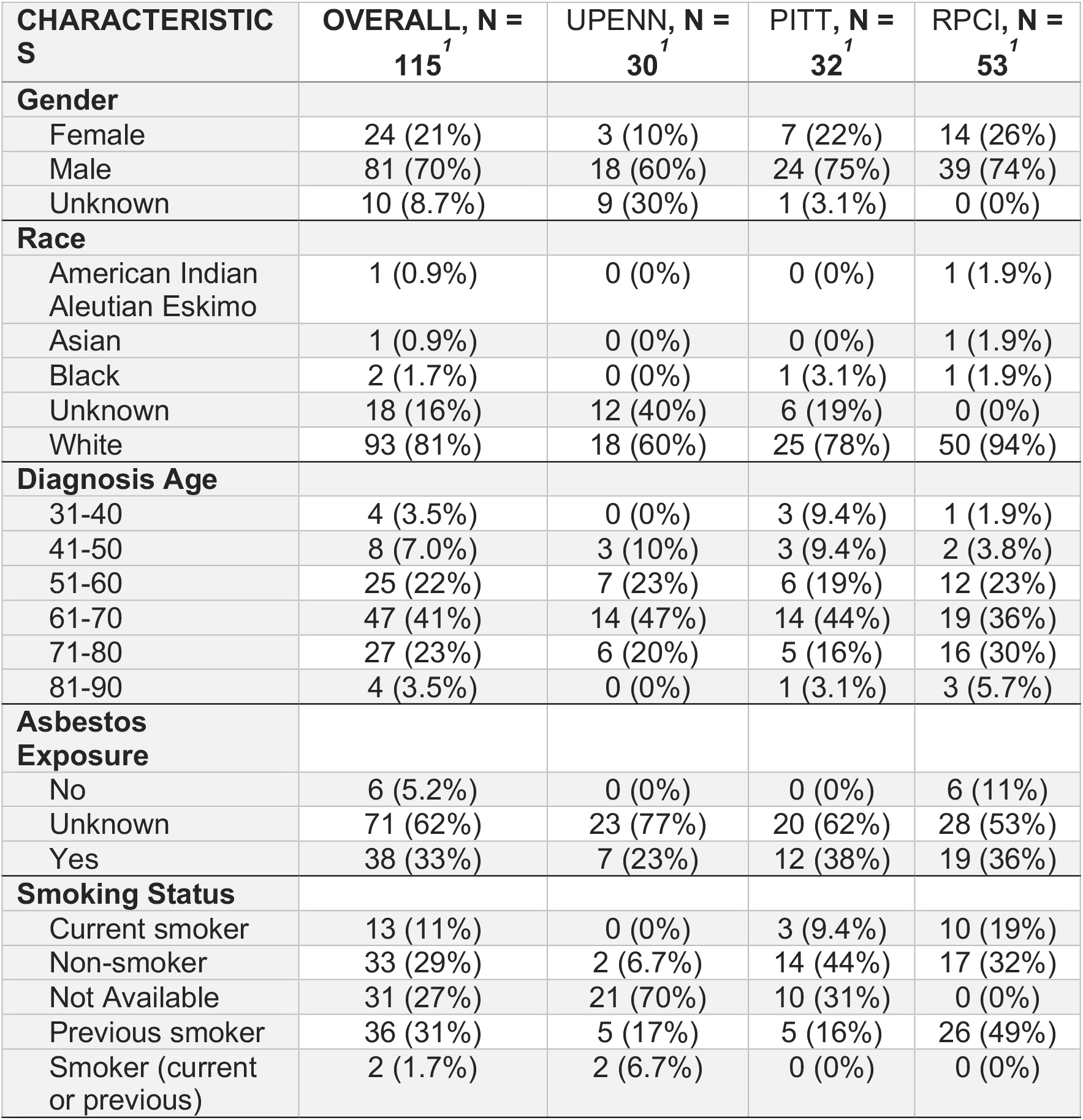
MM patient characteristics.

| **CHARACTERISTICS** | **OVERALL, N =**<br>**115<sup>1</sup>** | **UPENN, N =**<br>**30<sup>1</sup>** | **PITT, N =**<br>**32<sup>1</sup>** | **RPCI, N =**<br>**53<sup>1</sup>** |
| --- | --- | --- | --- | --- |
| **Gender** |  |  |  |  |
| Female | 24 (21%) | 3 (10%) | 7 (22%) | 14 (26%) |
| Male | 81 (70%) | 18 (60%) | 24 (75%) | 39 (74%) |
| Unknown | 10 (8.7%) | 9 (30%) | 1 (3.1%) | 0 (0%) |
| **Race** |  |  |  |  |
| American Indian<br>Aleutian Eskimo | 1 (0.9%) | 0 (0%) | 0 (0%) | 1 (1.9%) |
| Asian | 1 (0.9%) | 0 (0%) | 0 (0%) | 1 (1.9%) |
| Black | 2 (1.7%) | 0 (0%) | 1 (3.1%) | 1 (1.9%) |
| Unknown | 18 (16%) | 12 (40%) | 6 (19%) | 0 (0%) |
| White | 93 (81%) | 18 (60%) | 25 (78%) | 50 (94%) |
| **Diagnosis Age** |  |  |  |  |
| 31-40 | 4 (3.5%) | 0 (0%) | 3 (9.4%) | 1 (1.9%) |
| 41-50 | 8 (7.0%) | 3 (10%) | 3 (9.4%) | 2 (3.8%) |
| 51-60 | 25 (22%) | 7 (23%) | 6 (19%) | 12 (23%) |
| 61-70 | 47 (41%) | 14 (47%) | 14 (44%) | 19 (36%) |
| 71-80 | 27 (23%) | 6 (20%) | 5 (16%) | 16 (30%) |
| 81-90 | 4 (3.5%) | 0 (0%) | 1 (3.1%) | 3 (5.7%) |
| **Asbestos**<br>**Exposure** |  |  |  |  |
| No | 6 (5.2%) | 0 (0%) | 0 (0%) | 6 (11%) |
| Unknown | 71 (62%) | 23 (77%) | 20 (62%) | 28 (53%) |
| Yes | 38 (33%) | 7 (23%) | 12 (38%) | 19 (36%) |
| **Smoking Status** |  |  |  |  |
| Current smoker | 13 (11%) | 0 (0%) | 3 (9.4%) | 10 (19%) |
| Non-smoker | 33 (29%) | 2 (6.7%) | 14 (44%) | 17 (32%) |
| Not Available | 31 (27%) | 21 (70%) | 10 (31%) | 0 (0%) |
| Previous smoker | 36 (31%) | 5 (17%) | 5 (16%) | 26 (49%) |
| Smoker (current<br>or previous) | 2 (1.7%) | 2 (6.7%) | 0 (0%) | 0 (0%) |

**Table 2:** MM patient clinicopathological characteristics.

|  | **Overall**<br>**(n = 115<sup>1</sup>)** | **UPENN**<br>**(n = 30<sup>1</sup>)** | **PITT**<br>**(n = 32<sup>1</sup>)** | **RPCI**<br>**(n = 53<sup>1</sup>)** |
| --- | --- | --- | --- | --- |
| **Case Type** |  |  |  |  |
| Other | 2 (1.7%) | 0 (0%) | 2 (6.2%) | 0 (0%) |
| Peritoneal | 25 (22%) | 3 (10%) | 10 (31%) | 12 (23%) |
| Pleural | 88 (77%) | 27 (90%) | 20 (62%) | 41 (77%) |
| **Histology** |  |  |  |  |
| Benign Fibrous | 2 (1.7%) | 0 (0%) | 2 (6.2%) | 0 (0%) |
| Biphasic | 22 (19%) | 4 (13%) | 8 (25%) | 10 (19%) |
| Desmoplastic | 1 (0.9%) | 0 (0%) | 1 (3.1%) | 0 (0%) |
| Epithelioid | 69 (60%) | 22 (73%) | 17 (53%) | 30 (57%) |
| Fibrocystic | 1 (0.9%) | 0 (0%) | 1 (3.1%) | 0 (0%) |
| Multicystic | 1 (0.9%) | 0 (0%) | 0 (0%) | 1 (1.9%) |
| Not Specified | 8 (7.0%) | 0 (0%) | 0 (0%) | 8 (15%) |
| Papillary | 3 (2.6%) | 0 (0%) | 2 (6.2%) | 1 (1.9%) |
| Sarcomatoid | 8 (7.0%) | 4 (13%) | 1 (3.1%) | 3 (5.7%) |
| **Grade** |  |  |  |  |
| High | 46 (40%) | 11 (37%) | 27 (84%) | 8 (15%) |
| Intermediate | 17 (15%) | 15 (50%) | 2 (6.2%) | 0 (0%) |
| Low | 8 (7.0%) | 4 (13%) | 3 (9.4%) | 1 (1.9%) |
| Not Specified | 44 (38%) | 0 (0%) | 0 (0%) | 44 (83%) |
| **Survival Period** | 12 (7, 24) | 12 (9, 16) | 7 (3, 24) | 15 (8, 29) |
| Unknown | 3 | 0 | 3 | 0 |
| <sup>1</sup> n (%); Median (IQR) |  |  |  |  |

The integrated cohort (n_patients_=115) was made up of 81 males (70%) and 24 females (21%), with the rest unlisted. Ethnically, the cohort was composed of 93 white patients (81%), 2 black (1.7%), 1 Asian and 1 American Indian (0.9%), and the rest were unknown. The median age at diagnosis was in the 61–70 years age group, which represented 47 patients (41%). Other ages at diagnosis were 31–40 (3.5%), 41–50 (7%), 51–60 (22%), 71–80 (23%) and 81–90 (3.5%). Asbestos exposure was positive in 38 cases (33%), negative in 6 (5%), and unknown in the remaining cases. Also, 51 patients (44%) had a history of smoking, 33 were non-smokers (29%), and data was unavailable for the remaining 27%.

Clinically, the cohort was comprised of 88 cases of MPM (77%), and 25 cases (22%) of MPeM, while the remaining were other types of MM. Histologically, tumors were mostly of epithelioid origin (60%). Other histology types were 22 biphasic (19%), 8 sarcomatoid (7%), 3 papillary (2.6%), 1 benign fibrous (1.7%), 1 desmoplastic, 1 fibrocystic, 1 multicystic (0.9%) and 8 patients (7%) had cancer of unspecified histology. Median survival from time of diagnosis was 12 months, with an inter-quartile range of 7-24 months. We also performed Kaplan-Meier (K-M) analysis to compare the overall survival times between MPM and MPeM. In general, MPeM patients showed a higher overall survival than MPM patients in our cohort (log-rank test, *P* = 0.005). Further, patients with no asbestos exposure showed a higher overall survival than patients with asbestos exposure in our cohort (log-rank test, *P* = 0.015).

### Immune cell phenotypes characterized in malignant MM

The tumor microenvironment is a complex ecosystem of spatially resolved cellular interactions between tumor, stromal and immune cells. To gain further insight into the spatial distribution of key immune cell populations in MPM and MPeM, we performed mIF on TMA sections of MM samples using two Opal seven-color immunohistochemistry (IHC) kit (Akoya Biosciences) panels (**Fig. 1A**). In the first panel, tissue sections were stained for tumor cells (Pan-CK^+^), B cells (CD20^+^), CD4^+^ and CD8^+^ T cells, Tregs (CD4^+^Foxp3^+^), and macrophages (CD68^+^). In the second panel, DC (CD11c^+^) and NK (CD56^+^) cells were detected along with proteins LAG3 as well as BAP1, NF2, and MTAP as a proxy for frequent MM-associated TSG protein products.

To facilitate comparison between MPM and MPeM across different histological and pathological groups, we first performed cell quantification. Correlation analysis revealed a significant agreement in the cell population profiles between cores from the same patient compared to cores from different patients (average Pearson correlation 0.7403 and 0.5178 within and between different patients, respectively; *P* = 0.5×10^−30^) (**Supplementary Fig. 1)**.

Major immune cell populations included lymphocytes (CD8^+^ and CD4^+^ T cells) and CD68^+^ macrophages, whereas B cells and Tregs (CD4^+^Foxp3^+^) were relatively infrequent (**Fig. 1B**). Assessing the overall immune composition of MPM and MPeM, we observed a slightly higher proportion of CD8^+^ T cells in MPM (13.81%) compared to MPeM (8.89%), whereas the relative contribution of tumor cells (Pan-CK^+^) (39.16%) was higher in MPeM (39.16%) than in MPM (31.04%) (**Fig. 1C-D**). Although some cell type proportions other than CD8^+^ T and Pan-CK^+^ cells appeared to be different, these differences did not reach statistical significance due to the high variability among patients within tumor types.

In addition to the proportions of cell types in the MM TIME, their spatial organization should provide insights into how various cell types and their interactions influence MM development, progression, and therapeutic response. Hence, we computed their pairwise cell contact scores for the first panel including the tumor cytokeratin marker (Pan-CK^+^) (**Fig. 2A**). While the overall distribution of immune cells within the TIME was similar between MPM and MPeM, we did observe increased immune cell – tumor cell contacts in MPM compared to MPeM including tumor cell – CD4^+^ T cells/ CD8^+^ T cells/ macrophages/ Tregs (false discovery rate (FDR) < 0.1, **Fig. 2B-C, Supplementary Table 1**). Such higher tumor cell – immune cell interactions could potentially have implications for the immune response against the MPM vs. MPeM and could be a factor in the different behaviors of these two types of MMs.

**Figure 2:**
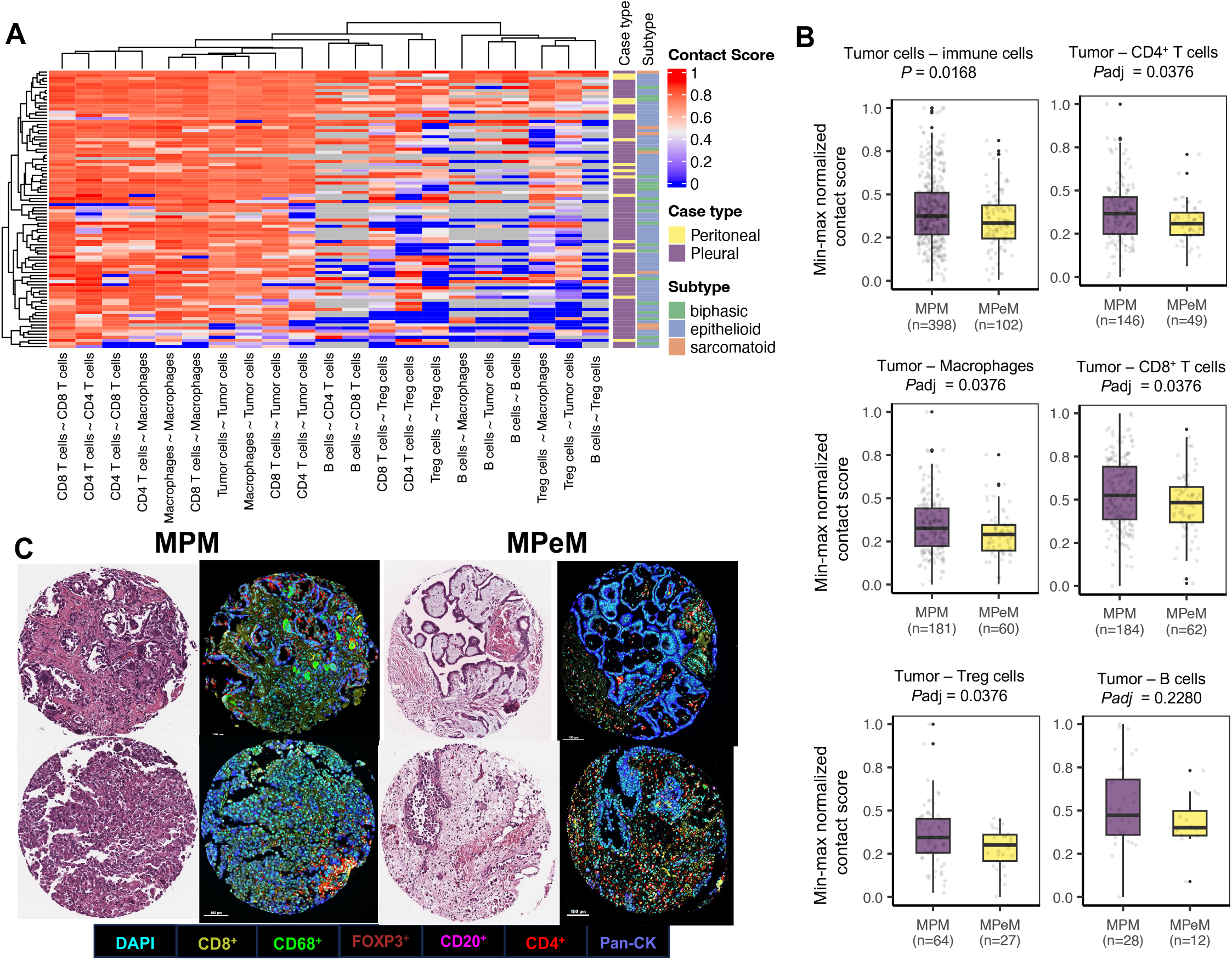
MM tumor immune cell contacts. **A**) Heatmap depicting cell-cell contacts in MPM and MPeM patients. Columns represent normalized pairwise cell-cell contacts, and rows represent samples. **B**) Tumor cell – overall immune cell contact scores as well as Tumor cell – CD4^+^ T cells/CD8^+^ T cells/macrophages/Treg contact scores in MPM versus MPeM patient cohorts. Statistics: FDR adjusted one-sided Multiple Mann–Whitney U test. Each dot represents the cell contact score of each core. **C)** Multiplex immunofluorescence (mIF), hematoxylin and eosin (H&E) staining, and cell type analyses in representative cases from two MPM and two MPeM. Images are shown at 10× magnification; scale bar = 100 μm.

### Immune cell phenotypes associated with clinical features and survival outcomes

We assessed the relationship between immune populations and clinical/pathological variables by interrogating the frequency of individual cell types as a percentage of total cells per patient. Each image was cross-referenced with clinical data from patients, including histology, survival, sex, age, asbestos exposure, smoking status, and tumor grade. Although the current classification of MM into epithelioid, sarcomatoid, and biphasic histology is prognostically useful, there remains significant variability in clinical features and patient outcomes within histologic subtypes. Indeed, immune cell proportions among histological groups were mostly similar (**Fig. 3A, Supplementary Table 2**). Epithelioid MPM tumors had higher proportions of B cells than biphasic ones, epithelioid and sarcomatoid MPM tumors had higher proportions of CD4^+^ T cells than biphasic tumors, and biphasic MPMs had higher amounts of Pan-CK^+^ cells than sarcomatoid tumors (*P* < 0.05, **Supplementary Table 2**). Although other cell type proportions appeared to be different, these differences did not reach statistical significance due to the high variability of cell frequencies among specimens. When we compared epithelioid/biphasic MPM and MPeM, we did not observe statistically significant differences in cell compositions (**Supplementary Table 3**).

**Figure 3:**
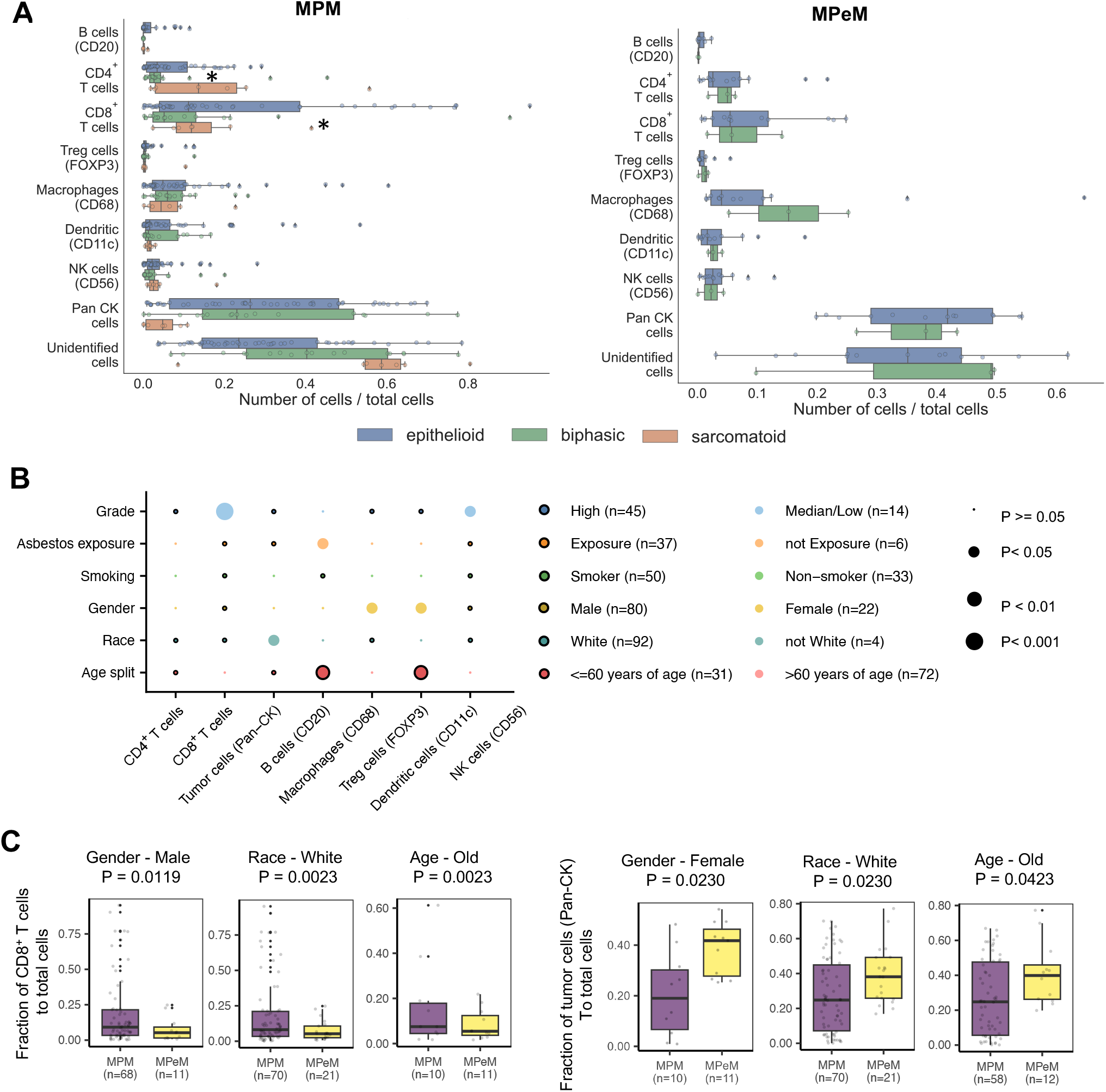
Immune cell phenotypes associated with clinical features. **A)** Prevalence of all cell types across MPM epithelioid (n=45), biphasic (n=16) and sarcomatoid (n=7) subtypes (left side) and MPeM epithelioid (n=12), and biphasic (n=3) subtypes (right-side) as a proportion of total cells. One-sided Multiple Mann–Whitney non-parametric U tests were used for statistical analysis, and *P* values of < 0.05 (*asterisks) were considered significant. **B)** Bubble plot in which the circle size represents the level of significance, and the circle color indicates which of the two comparisons on the *y-axis* has higher levels of the cell type on the *x-axis*. **C)** Box plots show cell type proportion across different clinic subgroups in MPM and MPeM. One-sided Multiple Mann–Whitney U test was used for statistical analysis, and P values < 0.05 were considered significant. The center of the box plot represents the median with 25th and 75th percentile bounds of the box and whiskers representing the least and highest value in the dataset.

We further performed exploratory survival analysis to determine if immune cell infiltration was associated with clinical outcome. We found that an increase in CD4^+^ T cells was associated with worse MPeM patient survival (log rank *P* = 0.0111) (**Supplementary Table 4**). Although we did not discover survival associations for several immune cell types, we found relationships between cell frequencies and specific clinical subgroups (**Fig. 3B**). We observed that median/low grade patients had higher proportions of CD8^+^ T cells and DCs compared to high grade patients. Patients without asbestos exposure had higher proportions of B cells. Female patients had higher proportions of macrophages and Tregs. Younger patients had higher proportions of FOXP3 and B cells. There were no significant associations between smoking and immune cell infiltration. When we compared MPM and MPeM patients based on gender, race and age, we observed statistically significant differences for CD8^+^ T and Pan-CK^+^ proportions (**Fig. 3C, Supplementary Table 5**). Together, these findings suggest that the spatial organization of TIME may provide additional insights into individual patient survival beyond histological subtype classifications and individual cell prevalence.

### MM architecture and TSG protein expression associations

To probe for the effect of BAP1, NF2, CDKN2A, and LAG3 expression at the protein level on MM TIME, we also stained TMAs with antibodies specific for BAP1, NF2, MTAP and LAG3. Multiple studies using a monoclonal anti-MTAP primary antibody report acceptable specificity and sensitivity of MTAP immunohistochemistry for the detection of *CDKN2A* homozygous deletion and diagnosis of MM^18^. In both MPM and MPeM, we observed that the average expression of the BAP1 protein was negatively correlated with the expression of CD8 and CD11c proteins (**Fig. 4A**) (Spearman’s correlation coefficient (rho) <= -0.4). CD8^+^ T cells (cytotoxic T cells) are a type of immune cell involved in directly attacking infected or cancerous cells, whereas DCs play a role in presenting antigens to other immune cells. Individual spot analyses revealed a significant positive correlation between tumor infiltration by NK cells (CD56^+^) and average BAP1, NF2, MTAP, and LAG3 expression for MPM (rho > 0.4) (**Fig. 4B**).

**Figure 4:**
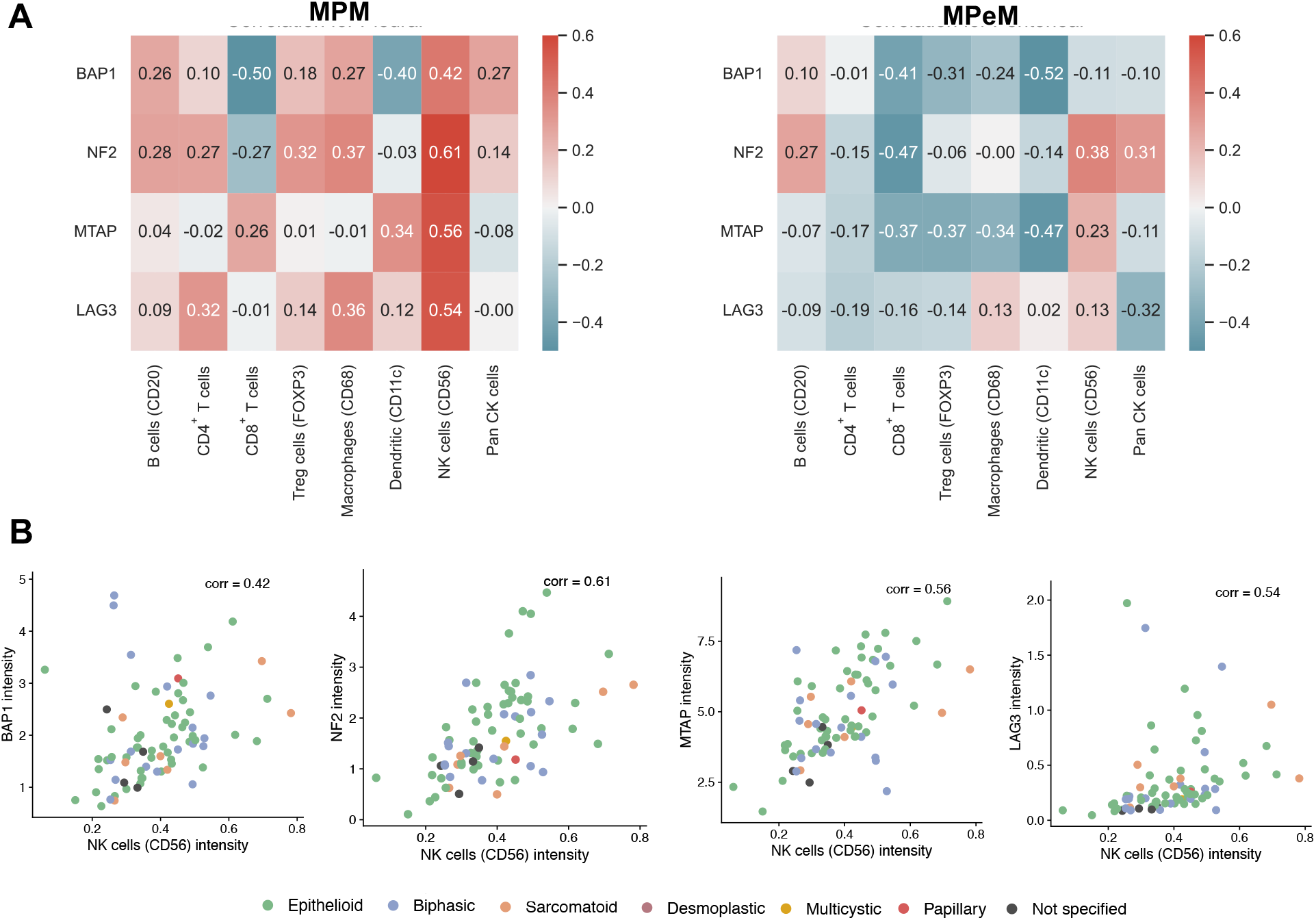
The association of BAP1, NF2, MTAP, and LAG3 intensity with immune cell type intensity in MPM and MPeM. Heatmap revealing correlations between mean intensity of immune cell markers and BAP1, NF2, MTAP, and LAG3 expression across biologically independent tumors in **A)** MPM and MPeM. **B)** Correlation of mean CD56 (NK cell) intensity and mean BAP1, NF2, MTAP, and LAG3. Dots are colored by histological subtypes as shown.

We observed a significantly greater tumor infiltration by (cytotoxic T cell) and CD11c (DC) was observed in MPM and MPeM tumors with low BAP1 expression (**Fig. 5A**). To further investigate the relationship between BAP1 and TIME, we computed pairwise contacts between different immune cell types and tumor cells from MM tumors with “BAP1-high” and “BAP1-low” expression levels (see Methods). Analysis of pairwise contacts in images of tumor samples revealed that the BAP1-low group was enriched with Pan-CK – CD8^+^ T cell contacts in MPM, but not in MPeM, cases (**Fig. 5B, C, Supplementary Table 6**). This finding could have implications for understanding the immune response and potential therapeutic strategies in patients with MPMs not retaining expression of BAP1 in tumor cells.

**Figure 5:**
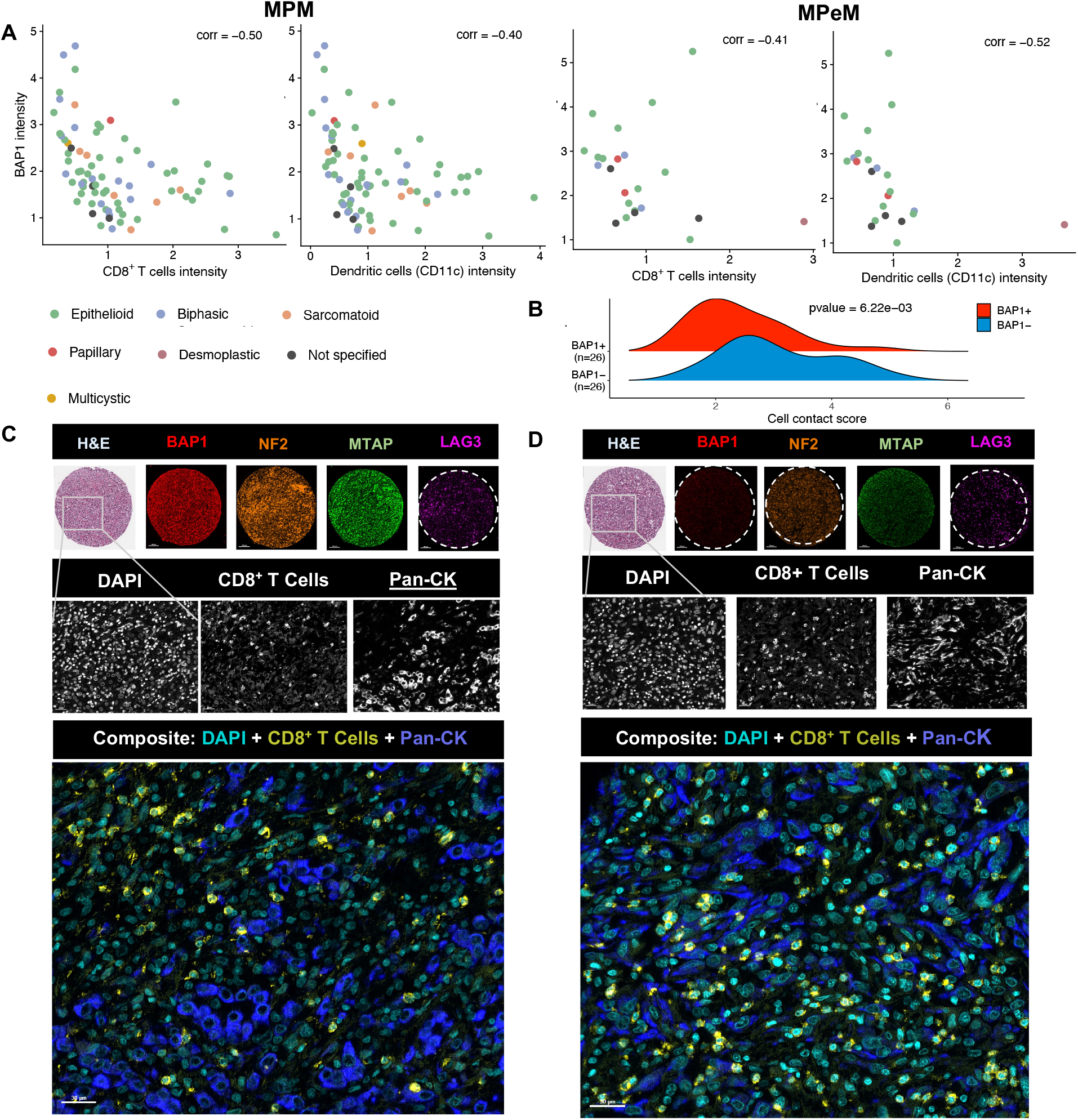
Analysis of cell types in the MPM BAP1-high and -low tumor groups. **A)** Correlation of mean CD8 (cytotoxic T cell) and CD11c (dendritic cell) intensities and mean BAP1, NF2, MTAP, and LAG3. Dots are colored by histological subtypes as specified in the key. **B)** Ridge density plot shows cell contact score between BAP1 and CD8^+^ T cells in the tumor region among BAP1 high and BAP1 low Pan-CK-enriched cores (CK percent > 20%). Note: density > 0.6 quantile and < 0.4 quantile are thresholds to separate cores as BAP1 high and BAP1 low. **C** and **D)** Segmented images showing increased interaction of cancer cells and CD8^+^ T cells in BAP1 low **(D)** versus BAP1 high MPM (**C**). *Upper panel:* TMA core sections stained with H&E, BAP1 – Opal 690, red; NF2 – Opal 780, orange; MTAP – Opal 620, green; and LAG3 – Opal570, magenta. Scale bars = 100 μm, 10× magnification. *Middle panel:* Zoomed in area from boxed segment in upper panel stained with DAPI – DNA (cyan); Opal 480 – CD8^+^ T cells (yellow); and Opal 780 – Pan-CK cells (blue). Scale bars = 10 μm, 100× magnification. *Lower panel:* Composite images. Scale bars = 30 μm, 30 × magnification.

### The Human Spatial Atlas of Malignant Mesothelioma

To support the research community in investigating our mIF MM dataset, we created the web resource called the Human Spatial Atlas of Malignant Mesothelioma (https://mesotheliomaspatialatlas.streamlit.app/). We provide interactive visualizations mIF and H&E images for each core in TMAs. Our web resource provides unique and valuable data for testing hypotheses about MM TIME that could facilitate the understanding of MM pathogenesis and identify novel therapies.

## Discussion

Here we applied mIF to MM TMAs and have provided a spatial map comparing the TIME of MPeM and MPM. This is the largest analysis performed on MM samples profiled by mIF to date. We also presented the Human Spatial Atlas of Malignant Mesothelioma (https://mesotheliomaspatialatlas.streamlit.app/), a web server containing mIF and H&E images.

Given the limited therapeutic options for patients diagnosed with MM, there is untapped translational value in understanding how the spatial architecture of the MM TIME relates to tumor biology, and whether specific immune cell subsets can be harnessed to improve clinical outcomes in this uniformly deadly disease. Our analysis has revealed T cells and macrophages to be significantly enriched in MM tumor tissues regardless of tumor site of origin. The MPE TIME generally had slightly more T cells, whereas the MPeM TIME had somewhat more macrophages. To explore the potential cross-talk between immune cells in MPM and MPeM, we used the cell-cell contact analysis approach and found that the TIME of these two tumors is largely similar except for the significantly more immune – tumor cell contacts in MPM tumors. Despite the relative abundance of T cells in these tumors, there have been only very modest responses to T cell-centric therapies alone such as CheckMate 743 study (NCT02899299) for MPM and MPeM. Indeed, Yin et al. recently showed high density and spatial proximity of CD8^+^ T cells to tumor cells were associated with better response to nivolumab, whereas the proximity of regulatory T cells to tumor cells was associated with worse response^19^. Our findings suggest that the distinct landscape of the TIME could be a potential predictor of the efficacy of immune checkpoint inhibitors (ICIs) in MPM.

Understanding how TSG expression relates to immune cell presence and function could have implications for immunotherapy strategies. If a tumor’s TSG status affects immune cell infiltration and activity, it might guide the development of treatments that modulate the tumor microenvironment to enhance T/NK/DC cell-mediated tumor clearance. Through our mIF analyses, we found that BAP1-low tumors have higher CD8^+^ T cell and DC infiltration. Thus, BAP1 expression could be used as a potential biomarker for T cell-centric therapies. However, our study delineated differences in observations between the two major types of MM, suggesting that MPM may be better at eliciting an adaptive immune response than MPeM. Interestingly, in MPeM, but not MPM, an inverse correlation was observed between the expression of BAP1 and the intensity of immune cell markers for CD8^+^ T cells, Tregs, macrophages, and DCs (**Fig. 4A**). For MPM, an inverse correlation was observed only between the expression of BAP1 and markers for CD8^+^ T cells and DC. This suggests that in MPeM, loss of BAP1 may correlate with tumor infiltration by multiple immune cell types, including CD8^+^ T cells, Tregs, macrophages, and DC, whereas in MPM, increased infiltration by CD8^+^ T cells and DC, but not other immune cells, is likely only when BAP1 alone is deleted. Whether such changes in the spatial landscape of the MM TIME would contribute, in part, to the generally more favorable prognosis of MPM than MPeM, or the better survival outlook of patients with alterations of *BAP1* alone among MPM cases, is not known but of future research interest.

A limitation of this study is that the analysis is based on staining small biopsy specimens in the TMA which may not be representative of the whole tumor. This is important as there is evidence of intra-tumor heterogeneity and even polyclonality in MM^20,21^. However, the majority of our tumors have multiple cores. The small sample size also limits statistical power and multivariable modeling flexibility. In addition, with only nine markers, we are restricted from distinguishing certain immune cell subtypes, such as exhausted versus effector CD8^+^ T cells or tumor-associated versus inflammatory macrophages, that are known to influence patient outcomes^9^. Another limitation of our study is the lack of matching clinical data about treatment allocation and response to treatment; this was not possible due to the retrospective nature of this analysis and limited available follow up data.

In conclusion, our study could help establish TIME architecture as a novel marker in MM. Our data represent a valuable resource (https://mesotheliomaspatialatlas.streamlit.app/), that adds to a quickly evolving body of literature supporting the importance of spatially resolved datasets in understanding how the TIME architecture relates to MM biology.

## Methods

### Multiplexed immunofluorescence (mIF)

The MM TMAs (n=3, comprising a total of 336 samples/cores of MM with predominantly epithelioid histology) from a total of 115 patients were obtained from NMVB. Consolidated de-identified patient data and tissue samples were also made available by NMVB. Multiplexed immunofluorescence of TMA was performed using the AkoyaBio manufacturer’s protocol. Antibody details are described in **Supplementary Table 7**. Two rounds of immunofluorescence staining were performed on sets of equivalent TMAs. The first round was performed using antibodies against pan-cytokeratin (epithelial cell marker), CD4 (helper T-cell marker), CD8 (cytotoxic T-cell marker), CD20 (B-cell marker), CD68 (macrophage marker), and FOXP3 (regulatory T-cell marker). Subsequently, a second round of immunofluorescence staining was performed with antibodies against CD11c (dendritic cell marker), CD56 (NK cell marker), BAP1, MTAP, LAG3, and NF2. In each round of staining, each fluorophore was linked to one antibody to detect the cellular target. Slides were baked in a dry oven at 60°C, de-paraffinized with xylene and ethanol, and finally re-fixed in 10% neutral buffered formalin for 15 minutes. Slides were stained according to manufacturer’s instructions for MoTIF Vectra panels (Akoya Biosciences). Microwave heat induced antigen retrieval (HIER) was performed, followed by blocking for 10 minutes. Primary antibodies were incubated for 30 minutes at room temperature. Secondary antibodies conjugated to horseradish peroxidase were then added for 10 minutes. Cells were stained with the following primary antibody/conjugated opal pairs: CD8/Opal480, CD4/Opal690, FOXP3/Opal570, CD68/Opal520, CD20/Opal620 and Pan-CK/Opal780 for the first panel. Cells were stained with the following primary antibody/conjugated opal pairs: CD11c/Opal480, BAP1/Opal690, LAG3/Opal570, CD56/Opal520, MTAP/Opal620, and NF2/Opal780 for the second set. Nuclei were stained with DAPI, and slides were mounted with coverslips using Pro-Long Diamond Anti-fade mounting media (ThermoFisher).

### Analysis of multispectral immunofluorescence images

TMA spots were imaged using Vectra® Polaris™ at 20× and analyzed using Akoya InForm Tissue Analysis Software and “phenoptr” which is an R script from Akoya (open source) that consolidates/analyses output tables generated in InForm. Within each TMA core, cells were automatically assigned cell types based on the marker expression using Akoya’s integrated inForm analysis software. Each cell was classified using the following surface markers: CD8^+^ for CD8^+^ T cells (cytotoxic T cells); CD4^+^FOXP3^−^ for CD4^+^ T cells; CD4^+^FoxP3^+^ for regulatory T cells (Tregs); pan-CK^+^ for tumor cells (CK); CD68^+^ for macrophages; CD11c^+^ for dendritic cells (DCs), CD56^+^ for Natural Killer cells (NKs); and CD20^+^ for B cells.

Measurement matrices consisting of centroid position (*x,y*), per-channel intensity, and cell types were exported and further processed in R (v4.0.3). For the first round of staining, cells that were positive for a single marker were assigned to the cell type corresponding to that marker as mentioned above. For cells with more than one positive marker, it was assigned to the marker with the highest intensity excluding Pan-CK. Cells without any positive marker were placed into a cell type designated “Unidentified”. Cell density was calculated using the following formula: the number of cells in the area divided by the whole tissue area (cells/mm^2^). Phenoptr and phenoptrReports were used for the identification of cellular subsets and spatial analysis including cell–cell distance. Distance between two cell subtypes was calculated using the x- and y-coordinates from the inForm raw data and further calculated by finding per-cell nearest neighbor distance using phenoptr. The distance between each cell and its nearest cell of both the same cell phenotype and the other cell phenotype were automatically calculated and the mean was determined.

### Cell type composition and density analysis

We had nine cell-type markers in two rounds of immunofluorescence staining in which CD56^+^ and CD11c^+^ were in the second panel and others are in the first panel. When generating combined cell type compositions, we subtracted CD56 and CD11c proportions (number of cells/total cells in the core) in the second panel from the “Unidentified” cell type proportions in the first panel to adjust the “Unidentified” value. Cell type proportion and density (number of cells/area) were calculated for each core (sample), then integrated to patient-wise proportion by averaging all cores belonging to the patient. Cell density values were log-transformed before statistical testing. To determine the difference in cellular distribution, the proportion of the cell types was compared across the type of MM and the histology types within the case types.

### Generation of cell-cell contact matrices

Cell-to-cell contact matrices were created using a custom-made Java algorithm with FCS files exported from Vortex, both of which were developed in Goltsev et al. ^22^. The direct neighbors of a cell were based on the Delaunay triangulations graph. Cell-to-cell contact score was the log odds ratio of observed adjacency frequency over the theoretical frequency of a pair of phenotypes, where the theoretical frequency was a simulation of beta distribution with the parameter derived from the cells and pair co-occurrence counts. The score value indicated the likelihood of the direct contacts of two phenotypes of the pair. The higher the score, the higher chance of contact. We performed a one-sided Mann–Whitney U test to compare groups. An FDR value <0.15 was considered statistically significant.

### Survival analysis

Kaplan–Meier curves and log-rank test were used for the analysis of overall survival association with cell type proportions and cell-cell contact scores. A P*-*value <0.05 was considered statistically significant. We designated the top 40% of the patients as the high-risk group and the bottom 40% as the low-risk group. The log-rank test was used to compare two Kaplan-Meier survival curves with the null hypothesis that there is no survival difference between the populations.

### Protein expression association analysis

When conducting an association between LAG3, BAP1, NF2, and MTAP protein expression and cell type marker expression, we used Spearman rank correlation to assess their relationship. For investigation on cell contacts between immune cells and Pan-CK associating with TSG protein level, BAP1 high and low patients were determined by >0.6 and <0.4 quantiles of the BAP1 density. Pan-CK proportion exceeding 20% was considered tumor-enriched core. Cell contact heatmap and the difference between MPM and MPeM analysis were performed on min-max normalized contact score.

### Statistical analysis and visualization

Statistical analyses of patient characteristics were performed using one-sided Mann–Whitney U test, and a significance was called when P-value < 0.05, a predetermined threshold. Graphs were generated using QuPath^23^, Phenochart™ 1.1 with MOTiF™ (Akoya Biosciences (R), RColor-Brewer (version: 1.1 2), ggplot2 (version: 3.4.2), gridExtra(version:2.3), viridis(version:0.6.4), ggpubr(version:0.6.0), ggtext(version:0.1.2) ComplexHeatmap (version: 2.16.0), ggrepel (version: 0.9.1), and ggridges(version:0.5.4) packages. For general data analysis and manipulation, stats (version:4.3.1), survival (version:35-5), survminer (version:0.4.9), coin (version:1.4-2), tidyverse (version:2.0.0), dplyr (version: 1.1.2), data.table (version: 1.14.8), and xlsx (version:0.6.5) were used with R (version 4.3.1) ; numpy (version:1.21.5), pandas (version:1.4.4), and scipy (version:1.7.3) were used with Python (version 3.9).

### Web portal for the database

We built a web portal to present user-friendly mIF and H&E image browser (https://mesotheliomaspatialatlas.streamlit.app/). All images can be searched, visualized, and downloaded from the web portal. The back end of the portal was implemented in Python (version: 3.11), while the front end was built through Streamlit (version: 1.25.0) open-source framework. The application was deployed to Streamlit cloud platform and is freely available (https://mesotheliomaspatialatlas.streamlit.app/) without registration or login. All the functions of the database have been tested in Google Chrome and Apple Safari browsers. Static figures and logos were made using BioRender. (www.biorender.com)

## AUTHOR CONTRIBUTIONS

HUO conceived the project, supervised the research/analysis and wrote the manuscript. XM performed the mIF analysis. DL organized clinical data. ESK prepared mIF and H&E figures. XM and ESK helped to write the manuscript. TB provided mIF data generation support. JRT engaged in supportive discussions during the course of the project and provided helpful suggestions and revisions of the manuscript. All authors have read and agreed to the published version of the manuscript.

## ACKNOWLEDGEMENTS

We would like to thank Koushul Ramjattun for his assistance in preparing conceptual figure as well as Sheryl Kunning and Marian Joy for performing multiplexed immunofluorescence. The authors acknowledge Sandhya Prabhakaran and Dario Vignali for the helpful discussion and suggestions. This work was funded by the National Institutes of Health (R35GM146989 and R00CA207871). ESK was supported by National Library of Medicine Training grant [T15 LM007059-24]. This work is supported by the Centers for Disease Control and Prevention (CDC) in association with the National Institute for Occupational Safety and Health (NIOSH) Grant, National Mesothelioma Virtual Bank or NMVB (U24OH009077). Data analyses in this research were supported by the University of Pittsburgh Center for Research Computing and the Extreme Science and Engineering Discovery Environment (XSEDE), which is supported by National Science Foundation grant number OCI-1053575. Specifically, it used the Bridges2 system, which is supported by NSF award number ACI-1445606, at the Pittsburgh Supercomputing Center.

## CONFLICT OF INTEREST STATEMENT

JRT has provided legal consultation regarding the role of genetic alterations in MM. The remaining authors declare no conflict of interest.

## DATA AVAILIBILITY

The multiplex IF and H&E images are available at https://figshare.com/projects/Mesothelioma_spatial_atlas/ and https://mesotheliomaspatialatlas.streamlit.app/. The remaining data are available within the Article, and Supplementary Information.

