## Supplemental Information for "Spatial landscape of malignant pleural and peritoneal mesothelioma tumor immune microenvironment"

**Supplementary Figures**

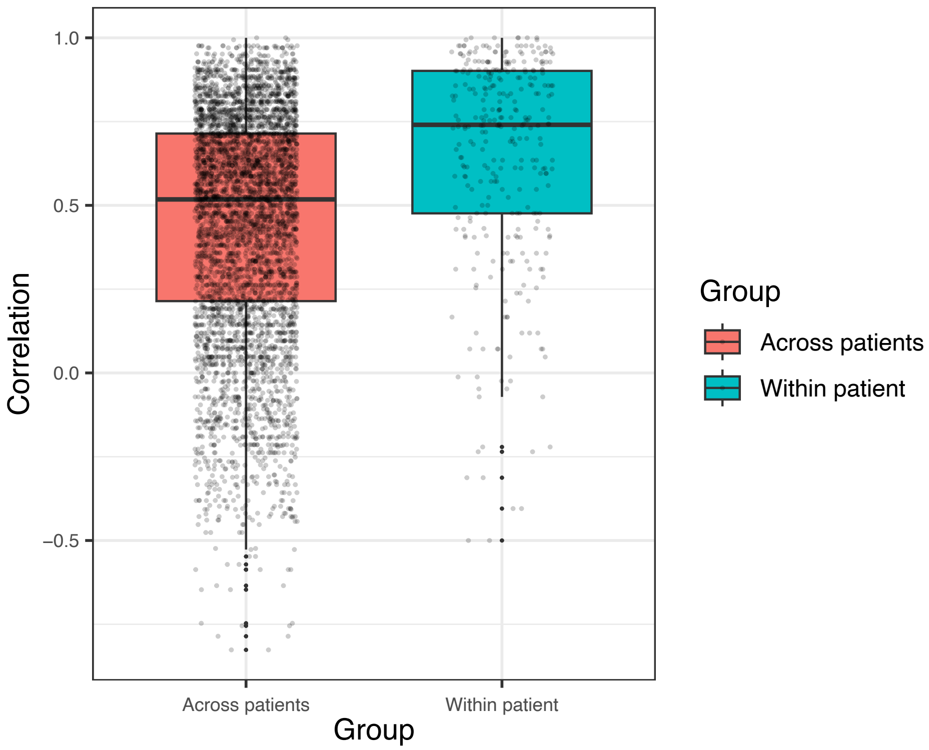


**Supplementary Figure 1:** TMA core cell-type proportions are more similar within patients than across patients. Each value in the boxplot represents the Pearson correlation coefficients between cell-type proportions across patients and within patients.

**Supplementary Tables**

**Supplementary Table 1: Comparison of cell-cell contact score differences between MPM and MPeM.** Two-sided Kruskal Wallis test was used for statistical analysis with FDR-adjusted <0.15 considered significant.

| **Contact pair** | ***P* value** | **FDR-adjusted** | **Mean contact score difference (MPM – MpeM)** |
|---|---|---|---|
| Tumor cells – tumor cells | 0.0002 | 0.0036 | 0.091 |
| $CD8^+$ T cells – $CD8^+$ T cells | 0.0012 | 0.0125 | -0.095 |
| $CD4^+$ T cells – tumor cells | 0.0112 | 0.0784 | 0.058 |
| Macrophages – tumor cells | 0.0158 | 0.0830 | 0.035 |
| Tregs – tumor cells | 0.0300 | 0.1054 | 0.041 |
| $CD8^+$ T cells – tumor cells | 0.0301 | 0.1054 | 0.043 |
| B cells – Macrophages | 0.0393 | 0.1118 | -0.214 |
| B cells – $CD8^+$ T cells | 0.0426 | 0.1118 | -0.051 |
| $CD8^+$ T cells – Macrophages | 0.0625 | 0.1458 | -0.069 |
| Macrophages – Macrophages | 0.0714 | 0.1499 | 0.026 |
| $CD8^+$ T cells – FOXP3 | 0.1020 | 0.1947 | -0.042 |
| $CD4^+$ T cells – $CD8^+$ T cells | 0.1180 | 0.2065 | -0.040 |
| B cells – B cells | 0.1620 | 0.2460 | -0.026 |
| B cells – $CD4^+$ T cells | 0.1640 | 0.2460 | -0.094 |
| B cells – tumor cells | 0.2280 | 0.3192 | 0.072 |
| Tregs – Tregs | 0.4020 | 0.5276 | 0.045 |
| $CD4^+$ T cells – Macrophages | 0.4550 | 0.5621 | -0.014 |
| $CD4^+$ T cells – Tregs | 0.5730 | 0.6540 | 0.014 |
| $CD4^+$ T cells – $CD4^+$ T cells | 0.5930 | 0.6540 | 0.022 |
| Tregs – Macrophages | 0.6240 | 0.6540 | 0.009 |
| B cells – Tregs | 0.6540 | 0.6540 | 0.097 |

**Supplementary Table 2: Distribution of immune/tumor cells across different histologies for MM cohort.** Table listing *P* values of individual immune/tumor cell subsets. Two-sided Kruskal Wallis test was used for statistical analysis with *P* < 0.05 considered significant.

| | **MPM** | | | **MPeM** |
|---|---|---|---|---|
| **Cell Type** | **Epithelioid vs. Biphasic** | **Epithelioid vs Sarcomatoid** | **Biphasic vs Sarcomatoid** | **Epithelioid vs. Biphasic** |
| B cells ($CD20^+$) | **0.044** | 0.095 | 0.416 | 0.217 |
| $CD4^+$ T cells | 0.371 | **0.037** | **0.040** | 0.473 |
| $CD8^+$ T cells | 0.053 | 0.475 | 0.077 | 0.420 |
| Tregs ($FOXP3^+$ $CD4^+$) | 0.163 | 0.256 | 0.447 | 0.469 |
| Macrophages ($CD68^+$) | 0.366 | 0.282 | 0.337 | 0.101 |
| DCs ($CD11c^+$) | 0.410 | 0.677 | 0.637 | 0.230 |
| NK ($CD56^+$) | 0.161 | 0.323 | 0.108 | 0.319 |
| Pan-$CK^+$ | 0.491 | **0.001** | **0.001** | 0.420 |

**Supplementary Table 3: Comparison of cell-type distributions between MPM and MPeM within same histologic subtypes***

| Cell Type | MPM vs. MPeM | |
|---|---|---|
| | **Epithelioid** | **Biphasic** |
| B cells (CD20$^+$) | 0.234 | 0.122 |
| CD4$^+$ T cells | 0.531 | 0.184 |
| CD8$^+$ T cells | 0.065 | 0.479 |
| Tregs (FOXP3$^+$ CD4$^+$) | 0.206 | 0.077 |
| Macrophages (CD68$^+$) | 0.625 | 0.093 |
| DCs (CD11c$^+$) | 0.689 | 0.246 |
| NK (CD56$^+$) | 0.342 | 0.521 |
| Pan-CK$^+$ | 0.072 | 0.184 |

*Table lists *P* values of differences for individual immune/tumor cell populations between MPM and MPeM tumors. One-sided Kruskal Wallis test was used for statistical analysis with $P < 0.05$ considered significant.

**Supplementary Table 4:** The association of immune cells proportion with overall survival in all MM as well as for MPM and MPeM separately

| | **MM** | **MPM** | **MPeM** |
|---|---|---|---|
| CD4$^+$ T cells | 0.17 | 0.67 | **0.02** |
| CD8+ T cells | 0.63 | 0.46 | 0.97 |
| Pan-CK+ | 0.20 | 0.10 | 0.93 |
| Tregs (FOXP3+ CD4+) | 0.13 | 0.38 | 0.39 |
| B cells (CD20+) | 0.27 | 0.38 | 0.35 |
| Macrophages (CD68+) | 0.12 | 0.33 | 0.12 |
| DCs (CD11c+) | 0.21 | 0.70 | 0.37 |
| NK (CD56+) | 0.36 | 0.72 | 0.36 |

*Table lists *P* values based on log-rank test. $P < 0.05$ considered significant.

**Supplementary Table 5:** Comparison of clinical parameters between MPM and MPeM for various cell types. FDR-corrected sided Wilcox test for sex, age, smoking status, asbestos exposure, stage

| | CD4+ T cells | CD8+ T cells | Pan-CK+ | B cells (CD20+) | Tregs (FOXP3+ CD4+) | Macrophages (CD68+) | DCs (CD11c+) | NK (CD56+) |
|---|---|---|---|---|---|---|---|---|
| High grade | 0.680 | 0.061 | 0.054 | 0.897 | 0.696 | 0.827 | 0.266 | 0.725 |
| Low/Median grade | NA | NA | NA | NA | NA | NA | NA | NA |
| Smoker | 0.876 | **0.131** | **0.071** | 0.897 | 0.696 | 0.592 | 0.515 | 0.722 |
| Non-smoker | 0.876 | **0.131** | 0.204 | 0.897 | 0.696 | 0.827 | 0.847 | 0.725 |
| Asbestos Exposure | **0.065** | **0.123** | **0.136** | 0.897 | 0.696 | 0.827 | 0.818 | 0.722 |
| No Asbestos Exposure | NA | NA | NA | NA | NA | NA | NA | NA |
| Young ( <60) | 0.876 | **0.091** | **0.122** | 0.897 | 0.721 | 0.592 | 0.815 | 0.725 |
| Old (>60) | 0.876 | **0.002** | **0.042** | 0.897 | 0.696 | 0.495 | **0.065** | **0.153** |
| White | 0.876 | **0.002** | **0.023** | 0.897 | 0.696 | 0.495 | **0.169** | 0.725 |
| Not Write | NA | NA | NA | NA | NA | NA | NA | NA |
| Male | 0.680 | **0.011** | **0.071** | 0.897 | 0.696 | 0.964 | 0.818 | 0.510 |
| Female | 0.680 | **0.131** | **0.023** | 0.897 | 0.696 | 0.592 | 0.266 | 0.153 |

FDR < 0.15 were bolded.

**Supplementary Table 6: Comparison of cell-cell contact score differences between MPM and MPeM***

| | **MPM** | | | **MPeM** | | |
|---|---|---|---|---|---|---|
| **Contact pair** | ***P* value** | **FDR-adjusted** | **$BAP1^{high}$ – $BAP1^{low}$** | ***P* value** | **FDR-adjusted** | **$BAP1^{high}$ – $BAP1^{low}$** |
| $CD4^{+}$ T cells – tumor cells | 0.074 | 0.120 | -0.685 | 0.549 | 0.549 | 0.226 |
| $CD8^{+}$ T cells – tumor cells | **0.003** | **0.013** | -0.521 | **0.046** | **0.093** | -0.630 |
| Macrophages – tumor cells | 0.090 | 0.120 | -0.147 | 0.036 | 0.093 | 0.756 |
| B cells – tumor cells | NA | NA | -0.147 | NA | NA | 0.756 |
| Tregs – tumor cells | 0.500 | 0.500 | -0.460 | 0.500 | 0.549 | -0.199 |

*Table lists *P* values and FDR values. Two-sided Kruskal Wallis test was used for statistical analysis with FDR-adjusted such that *P* < 0.15 was considered significant.

**Supplementary Table 7:** Antibody details

| Antibody | Clone | Concentration | Supplier | Cat. # |
|---|---|---|---|---|
| FOXP3 | D2W8E | 1:200 | Cell Signaling | 98377S |
| CD8 | C8/144B | 1:150 | Biocare Medical | ACI3160A |
| CD68 | D4B96 | 1:800 | Cell Signaling | 76437S |
| Pan-cytokeratin | AE1/AE3 | 1:150 | Santa Cruz Biotechnologies | sc-81714 |
| CD20 | L26 | 1:300 | Leica Biosystems | NCL-L-CD20-L26 |
| CD4 | EP204 | pre-dilute | Biocare Medical | API3209AA |
| LAG3 | D2G4O™ | 1:250 | Cell Signaling | 45208S |
| CD11c | D3V1E | 1:400 | Cell Signaling | 45581S |
| CD56 | BC56C04 | 1:100 | Biocare Medical | CM164A |
| BAP1 | N/A | 1:100 | Biocare Medical | ACI3247A |
| MTAP | N/A | 1:500 | Abcam | Ab96231 |
| NF2 | N/A | 1:300 | Sigma | HPA003097 |
